# A Rad51-Independent Pathway Promotes Single-Strand Template Repair in Gene Editing

**DOI:** 10.1101/2020.02.24.962423

**Authors:** Danielle N. Gallagher, Nhung Pham, Annie M. Tsai, Abigail N. Janto, Jihyun Choi, Grzegorz Ira, James E. Haber

## Abstract

The Rad51/RecA family of recombinases perform a critical function in typical repair of double-strand breaks (DSBs): strand invasion of a resected DSB end into a homologous double-stranded DNA (dsDNA) template sequence to facilitate repair. However, repair of a DSB using single stranded DNA (ssDNA) as a template, a common method of CRISPR/Cas9-mediated gene editing, is Rad51-independent. We have analyzed the genetic requirements for these Rad51-independent events in *Saccharomyces cerevisiae* in two different assays. Gene editing events were carried out either by creating a DSB with the site-specific HO endonuclease and repairing the DSB with 80-nt single-stranded oligonucleotides (ssODNs) or by using a bacterial retron system that produces ssDNA templates *in vivo* in combination with DSBs created by Cas9. We show that single strand template repair (SSTR), is dependent on Rad52, Rad59, Srs2 and the Mre11-Rad50-Xrs2 (MRX) complex, but not Rad51, Rad54 or Rad55. Srs2 acts to prevent overloading of Rad51 on the ssDNA filament, whereas Rad59 appears to alleviate the inhibition of Rad51 on Rad52’s strand annealing activity; thus, deletion of *RAD51* suppresses both the *srs2*Δ and *rad59*Δ phenotypes. This same suppression by *rad51*Δ of *rad59*Δ is found in another DSB repair pathway, single strand annealing (SSA). In contrast, gene targeting using an 80-bp dsDNA template of the same sequence is Rad51-dependent. We also examined SSTR events in which the ssODN carried several mismatches. In the absence of the mismatch repair protein, Msh2, we found that the fate of mismatches carried on the ssDNA template are very different at the 3’ end, which can anneal directly to the resected DSB end, compared to the 5’ end. We also find that DNA polymerase Polδ’s 3’ to 5’ proofreading activity frequently excises a mismatch close to the 3’ end of the template, similar to its removal of heterologies close to the 3’ invading end of the DSB. We further report that SSTR is accompanied by a 600-fold increase in mutations in a region adjacent to the sequences directly undergoing repair. These DNA polymerase ζ-dependent mutations may compromise the accuracy of gene editing.

## Introduction

DSBs are repaired through one of two pathways: homologous recombination (HR) or nonhomologous end joining (NHEJ). Both classical NHEJ and microhomology-mediated end joining (MMEJ) involve DNA ligase-mediated joining of the broken chromosome ends, which usually results in small insertions or deletions (indels) at the junction [1,2,3,4,5,6]. HR is a less mutagenic form of DSB repair, as it makes use of a homologous sequence as a donor template for repair. The template can be located on a sister chromatid, a homologous chromosome, or at an ectopic site. The majority of HR events are dependent on a core group of proteins, including the Rad51 strand-exchange protein that is responsible for homology recognition and initiating strand invasion into a double-stranded DNA (dsDNA) template [7]. In budding yeast, Rad51 interacts with and is assisted by numerous proteins, most significant being the mediators Rad52 and the Rad51 paralogs, Rad55 and Rad57, as well as the chromatin remodeler, Rad54 [8,9,10,11,12,13]. Rad52 also plays a key role in later steps of DSB repair, facilitating second-end capture of the DNA polymerase-extended repair intermediate [14].

However, some DSB repair events, though still requiring Rad52, are Rad51-independent. The best studied mechanism is single-strand annealing (SSA), where homologous sequences flanking a DSB are rendered single-stranded by 5’ to 3’ exonucleases and then annealed, creating genomic deletions [15,16,17,18]. SSA requires the Rad52 paralog Rad59, especially when the size of the flanking homologous regions is small [19]. A second Rad51-independent mechanism involves break-induced replication (BIR) [20, 21]. Rad51-independent BIR is also independent of Rad54, Rad55, and Rad57; however, Rad59 and a paralog of Rad54 called Rdh54/Tid1 assume important roles. Rad51-independent BIR also requires the Mre11-Rad50-Xrs2 complex, whereas Rad51-mediated events and SSA are merely delayed by the absence of these proteins [18]. A third Rad51-independent pathway, another form of BIR, operates to maintain telomeres in the absence of telomerase (known as Type II events). Here too, Rad59 and the MRX complex, as well as Rad52, are necessary, whereas the Type I Rad51-dependent telomere maintenance pathway does not require either Rad59 or MRX [22,23,24]. Similarly, DSB repair by intramolecular gene conversion involving short (33-bp) regions of homology is inhibited by Rad51 and is dependent on the MRX complex, Rad59, and Rdh54 [25]. Rad51-independent BIR pathways are also dependent on the Srs2 helicase that antagonizes loading of Rad51 onto resected DSB ends [21, 26].

Use of the RNA-guided CRISPR/Cas9 endonuclease has revolutionized gene editing in eukaryotic systems ranging from yeast to mammals [27,28,29]. Guided endonucleases are programmed to create site-specific DSBs that can be repaired by providing a homologous template [30, 31]. One approach that has been shown to be an efficient method of gene editing in a variety of eukaryotic systems is to introduce short single-stranded oligo deoxynucelotides (ssODNs) as a donor template [32,33,34,35].

Here we have examined the genetic requirements for single strand template repair (SSTR) in budding yeast, using two different systems: 1) an inducible HO endonuclease and an 80-nucleotide (nt) ssODN as a template for DSB repair, and 2) an optimized bacterial retron system to produce ssDNA templates *in vivo* with a targeted Cas9-mediated DSB. We confirm that in budding yeast, as in other eukaryotes, SSTR is a Rad51-independent mechanism, but show that this pathway is distinct from the previously-described Rad51-independent recombination pathways. SSTR depends on Rad52, Rad59, and Srs2 proteins, as well as the MRX complex, but is independent of Rdh54/Tid1. Surprisingly, deleting both Rad51 and Rad59 or deleting both Rad51 and Rdh54 leads to a significant increase in SSTR over WT or *rad51*Δ levels. We show a similar suppression of *rad59*Δ by *rad51*Δ in SSA. We conclude that Rad59 prevents Rad51 from inhibiting Rad52-mediated strand annealing. In contrast, this novel form of repair is specific to ssDNA, as dsDNA templates of the same size and sequences use a canonical Rad51-dependent process.

Additionally, by analyzing the fate of mismatches between the ssODN and the target DNA, we show that the fate of mismatches at the 5’ and 3’ ends of the template are differently incorporated into the gene-edited product. We show that both the *MSH2* mismatch repair (MMR) protein and the 3’ to 5’ exonuclease activity of DNA Pol*ζ* play important roles in resolving heteroduplex DNA. Finally, we demonstrate that SSTR is accompanied by a 600-fold increase in mutations in the region adjacent to the site of gene editing, which is dependent on the error-prone DNA polymerase *ζ* that fills in single-stranded regions generated during DSB repair.

## Results

### Single Stranded Template Repair is Rad51-Independent

As a model for DSB-induced gene editing, we used a galactose-inducible HO endonuclease to create a site-specific DNA break at the *MATα* locus of chromosome 3. In this strain, both *HML* and *HMR* donors have been deleted, so that a DSB that can only be repaired via NHEJ unless an ectopic donor is provided [1]. When HO is continually expressed, imprecise NHEJ repair occurs in approximately 2 x 10^-3^ cells, distinguishable by indels in the cleavage site that prevent further HO activity. Repair by homologous recombination can be accomplished by introducing an 80-nt ssODN as a repair template. Cells were transformed with an ssODN template and then plated onto media containing galactose, which rapidly induces an HO-mediated DSB [36]. The template contains 37-nt of perfect homology to each end of the DSB, surrounding a 6-nt *Xho*I restriction site (Fig. 1A). SSTR leads to the disruption of the HO cleavage site by the insertion of the *Xho*I site, whose presence can be confirmed by an *Xho*I digest of a PCR product spanning the region (Fig. S1). In WT cells, we achieve an editing efficiency of 75-90% among in DSB survivors, with the remaining survivors repaired via NHEJ; these indels are eliminated by mutants such as *mre11*Δ, *rad50*Δ, and *yku70*Δ that are known to be required for NHEJ (Fig. 1B). It should be noted that cell survival in this assay is very low, with an average survival rate of 2.8% (Fig. S2); this low efficiency reflects limitations in transforming ssODN, as shown below.

**Figure 1.**
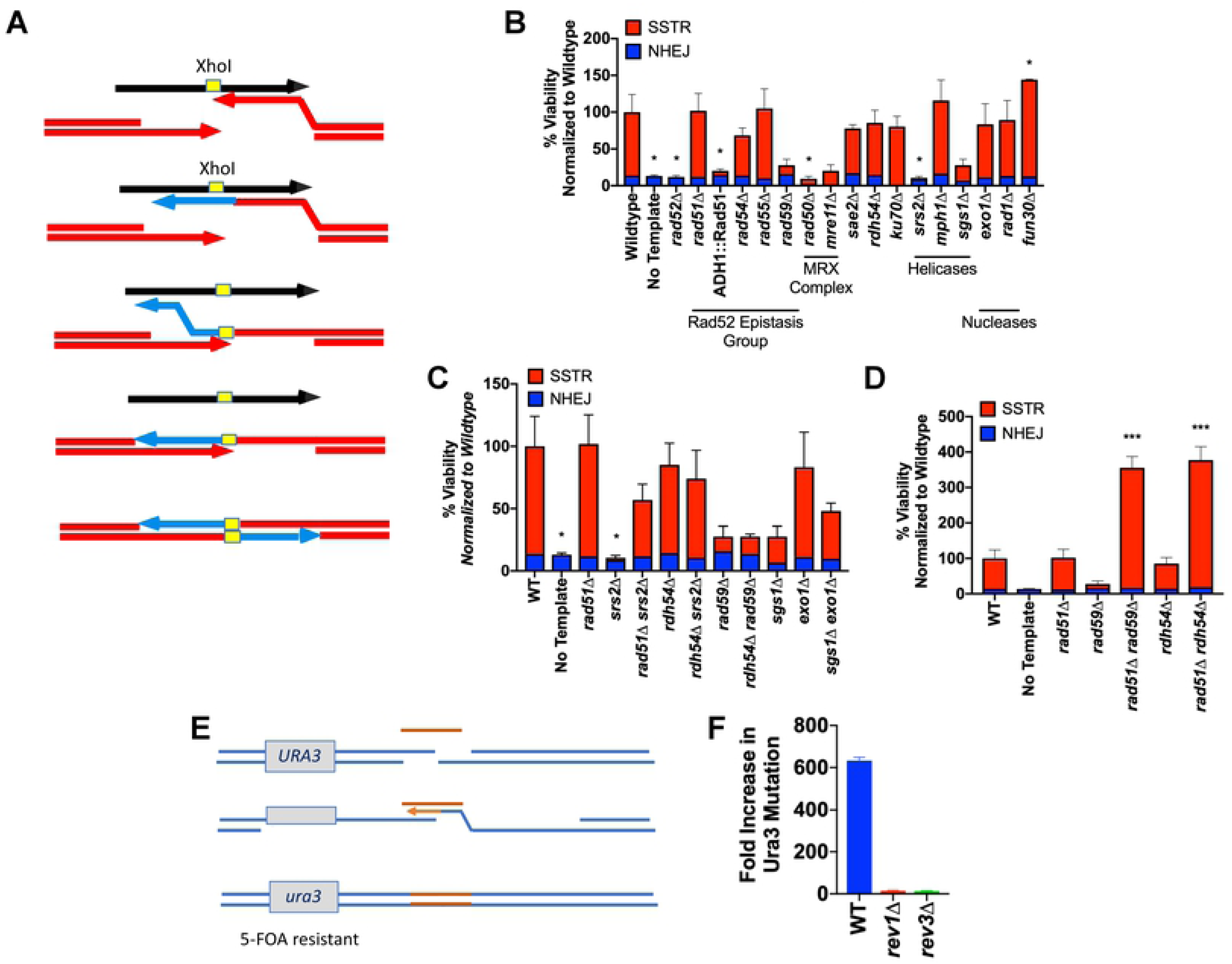
SSTR is a novel form of Rad51-independent DSB repair. A) A DSB was created at the *MATα* locus (red arrow) via induction of a galactose-inducible HO endonuclease. A ssODN with 37-nt homology on either side of an *Xho*I site (yellow) provides a template for SSTR, resulting in the insertion of an *Xho*I restriction site into the *MAT* locus. B) Viability determined by plate counts from galactose-containing media (induction media) over YEPD (non-induction media). Proportion of SSTR (red) and NHEJ (blue) events in various mutants involved in DSB repair are determined via PCR of the MAT locus, followed by an *Xho*I digest (Fig. S1). C and D) Effect of double mutants on SSTR. E) Effect of SSTR on a *URA3* gene integrated 200 bp upstream of the HO cleavage site. Mutations in *URA3* following SSTR were collected by replica plating survivors on galactose media onto 5-FOA medium (F). The increase in mutation rate in SSTR over the spontaneous mutation rate (determined by a fluctuation analysis) is eliminated by deleting components of Pol*ζ*. Significance determined using a paired t-test with Welch’s correction. * p ≤ 0.01, *** p ≤ 0.0001

We applied this assay to determine which recombination factors are required for SSTR. Consistent with previous results in both budding yeast and metazoans [34,35,37,38], SSTR proved to be Rad51-independent. Furthermore, SSTR was significantly inhibited when Rad51 was overexpressed from an *ADH1* promoter on a multicopy 2*μ* plasmid. SSTR also proved to be independent of the Rad51 paralog, Rad55, and the Rad54 translocase/chromatin remodeler, both of which are required for most DSB repair events that involve dsDNA templates. As expected, SSTR was dependent on the single strand annealing protein Rad52, with essentially all survivors resulting from NHEJ events. There was also a less profound reduction of SSTR in the absence of the Rad52 paralog Rad59 (Fig. 1B). Although in this instance the effect of *rad59*Δ was not statistically significant, in subsequent assays shown below Rad59 proved to be required for SSTR.

In previous studies of Rad51-independent recombination, both Rdh54 and the MRX complex were required [21, 25]. For SSTR, while MRX is required, Rdh54 is not, distinguishing SSTR from previously studied Rad51-independent pathways. It is also notable that while the MRX complex is required, Sae2 – which often functions in conjunction with MRX [39] – is not.

SSTR also requires the Srs2 helicase (Fig. 1B). One major function of the Srs2 helicase is to act as an anti-recombination factor by stripping Rad51 from the ssDNA tails formed after resection [40, 41]; thus, *srs2*Δ might mimic the inhibition of SSTR that is seen when Rad51 is overexpressed. Indeed, deleting *RAD51* suppressed *srs2*Δ’s defect in SSTR (Fig. 1C). Moreover, deleting *RDH54* suppressed the defect in *srs2*Δ. Although these results might suggest that Rdh54 acts in the same pathway as Rad51, their mutations are not epistatic; instead, deletion of both Rad51 and Rdh54 results in a 7-fold increase in SSTR over the levels seen in WT or in either of the single deletions (Fig. 1D).

Unexpectedly, deletion of Rad51 bypasses the requirement of Rad59 in SSTR and results in a much higher level of gene editing than simply deleting Rad51 (Fig. 1D). To further examine the genetic interaction between Rad51 and Rad59, we turned to a well-characterized SSA system. Previous studies have shown that Rad59 is important in SSA, especially when the length of the flanking homologous repeats is small [19]. Here, an HO-induced DSB within a *leu2* gene is repaired by SSA with a direct *leu2* repeat located 5 kb away, resulting in a chromosomal deletion of the sequences located between the two repeats, as well as one of the partial copies of the *leu2* gene (Fig. 2A). SSA is severely impaired in a *rad52-R70A* separation-of-function mutant that is capable of loading Rad51 onto ssDNA, but is defective in strand annealing [42] (Fig. 2B). Deletion of Rad59 has a similar inhibitory effect on SSA, lowering the efficiency of SSA to approximately 30% [19] (Fig. 2C). Deletion of Rad51 by itself only has a mild negative effect on SSA. Interestingly, deletion of Rad51 partially suppressed the SSA deficiency of Rad59 mutant strains (Fig. 2C). This suppression was not observed in either *rad52-R70A or rad52-R70A rad59*Δ, suggesting that Rad52-mediated strand annealing is increased in the *rad59*Δ *rad51*Δ double mutant. We propose that Rad59 alleviates the inhibition of Rad51 on the annealing activity of Rad52, freeing up the resected ssDNA tails for Rad52 to bind and perform its function as a strand annealing protein [43]. Since deletion of Rad51 in combination with either Rad59 or Rdh54 had a marked increase on the rate of SSTR, we also asked if a *rdh54*Δ *rad59*Δ might show a similar increase in SSTR; however, *rad59*Δ is not suppressed by *rdh54*Δ (Fig. 1D).

**Figure 2.**
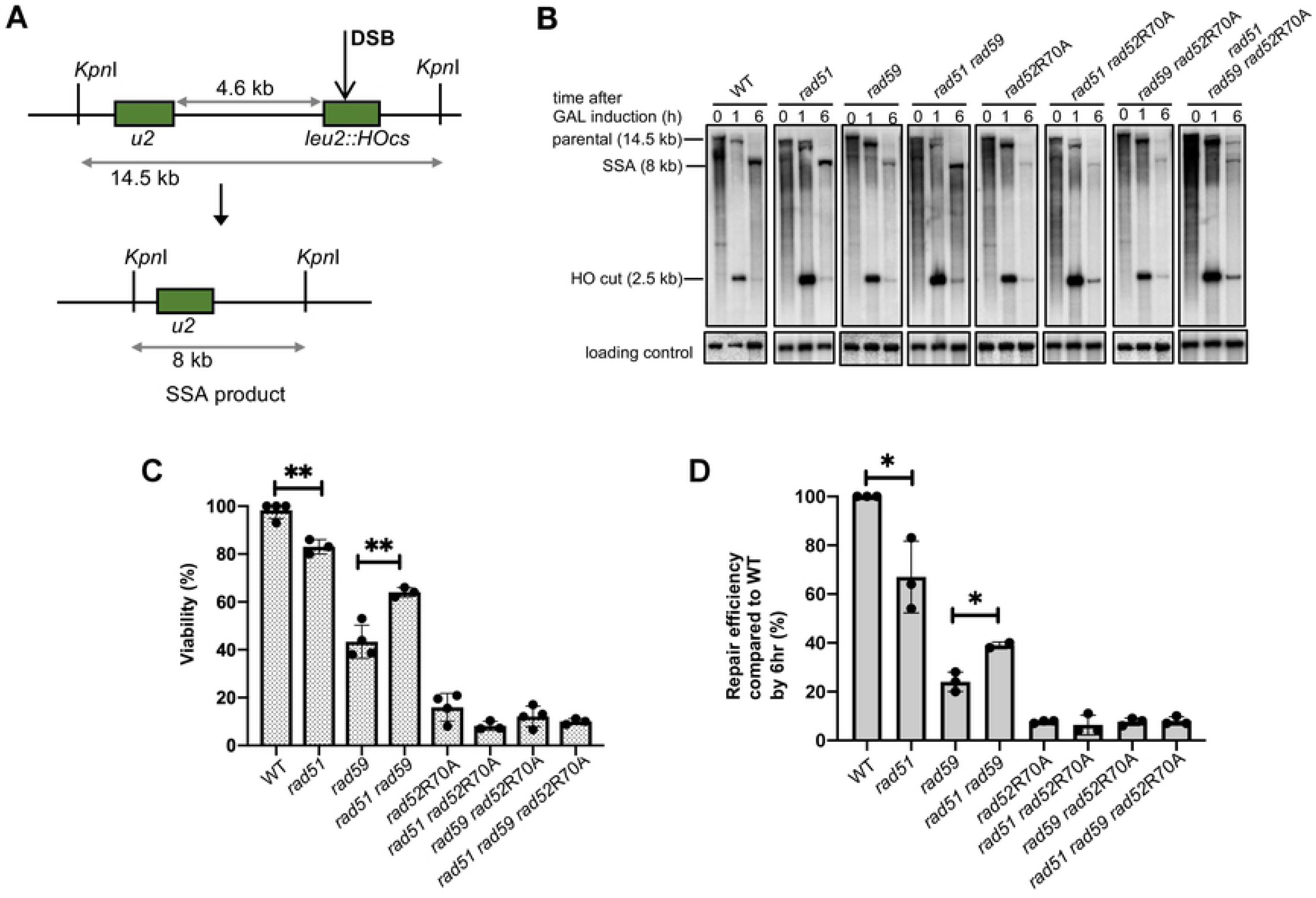
Rad59 alleviates the inhibition of Rad51 on annealing activity of Rad52. A) An HO-induced DSB results in repair via SSA between partial *leu2* gene repeats located 5 kb apart on the left arm of chromosome 3. B) Representative southern blots showing DSB repair products by SSA in WT and indicated mutants. C) Viability of mutants on galactose-containing plates, where HO DSBs are repaired via SSA (mean ± SD; n=3). Welch’s t-test was used to determine the p-value. D) Graphs show quantitative densitometric analysis of repair efficiency by 6 hr compared to WT (mean ± SD; n=3). Welch’s t-test was used to determine the p-value.

We also examined the role of several genes that are involved in the 5’ to 3’ resection of DSB ends: the Exo1 exonuclease and the Sgs1-Rmi1-Top3-Dna2 helicase/endonuclease complex [44, 45]. Deleting either Sgs1 or Exo1 had no effect on the efficiency of *Xho*I insertion, and neither did the *sgs1*Δ *exo1*Δ double mutant (Fig. 1D). These results suggest that the MRX complex can provide sufficient end resection to allow SSTR involving the homologous 37 nt of the ssODN donor. Although deleting the Fun30 SWI/SNF chromatin remodeler has been shown to strongly retard 5’ to 3’ resection of DSB ends [46,47,48], we found a significant increase in SSTR. Previously, deleting Fun30 was shown to protect double-stranded DNA fragments used in “ends-out” transformation [46], but in the present scenario, the transformed DNA is single-stranded. Whether Fun30 also affects the stability of ssODN’s is not clear. It should also be noted that in previous research using human cancer lines, SSTR was dependent on proteins in the Fanconi anemia (FA) pathway [49]. The helicase function of Mph1 is the only homolog of the FA pathway found in yeast, but Mph1 does not appear to play a role in SSTR in our system (Fig. 1B).

### SSTR is Accompanied by Frequent Mutations in Adjacent Regions

Previous research has suggested that DSB repair that involves DNA resection is highly mutagenic because ssDNA regions created during resection must be filled in once HR has completed [50,51,52]. Gap-filling, either by DNA polymerase *δ* or by translesion DNA polymerases such as Pol*ζ* have been shown to be responsible for mutation rates 1000-fold over background spontaneous mutation rates [53, 54]. Since SSTR likely involves extensive end resection and gap-filling, it is possible that there are significant off-target effects that have not previously been considered. To examine this possibility, the yeast *URA3* gene was inserted 200 bp centromere-proximal to the HO cleavage site and thus beyond the 37 nt of homology shared between the DSB end and the ssODN template. These cells were then targeted in the same *Xho*I insertion assay previously described (Fig. 1E). SSTR survivors were collected and then replica-plated onto 5-fluroorotic acid media (FOA), which selects for *ura3* mutants [55]. Compared to the spontaneous mutation rate, determined by fluctuation analysis [56], there was a >600-fold increase in *ura3* mutants after gene editing events (Fig. 1F). After deleting Rev1 or Rev3, components of Pol*ζ*, very few *ura3* mutants were recovered, indicating that the error-prone Pol*ζ* is responsible for these mutagenic events during SSTR (Fig. 1F). However, deletion of neither Rev1 nor Rev3 affects cell viability, indicating that other, less mutagenic mechanisms can be used for gap-filling in the absence of Pol*ζ* (Fig. S3). These data suggest that adjacent off-target effects of gene editing pose a danger that should be ruled out in selecting gene-editing events.

### Genetic Requirements of SSTR are Identical Using Cas9

Although it was convenient to survey many mutations using the highly efficient and easily inducible HO endonuclease, we confirmed that the same genetic requirements apply when a DSB is created by CRISPR/Cas9. Given the low efficiency of transforming ssODNs into yeast and screening for Cas9-mediated gene editing events, we turned to a modified version of the CRISPEY system to produce ssDNA templates *in vivo* [57]. This system utilizes a yeast-optimized *E. coli* retron system, Ec86, to generate designer ssDNA sequences *in vivo*. Retrons are natural DNA elements encoding for a reverse transcriptase (RT) that acts on a specific template sequence to generate single stranded DNA products [58,59,60]. These ssDNA products are covalently tethered to their template RNA by the RT, however after reverse transcription the RNA template is degraded [61, 62]. The CRISPEY system utilizes a chimeric RNA of Ec86 joined to the gRNA scaffolding of Cas9 at the 3’ end [57]. By integrating a yeast-optimized galactose-inducible Cas9 and retron (RT) onto chromosome 15, and using a CEN/ARS plasmid containing a galactose-inducible gRNA linked to the retron donor template, we were able to achieve high efficiency of gene editing at two different chromosomal locations, within the *MAT* locus near the HO cleavage site, and at a 5-bp insertion in the *lys5* locus (Fig. 3A and 3B). Successful SSTR via an 80-nt retron-generated ssDNA donor carrying the wild type *LYS5* sequence results in Lys^+^ recombinants that are easily recovered by replica plating galactose-induction survivors onto lysine drop out media. Compared to the <3% of cells that properly inserted the *Xho*I site at *MAT* when ssDNA was transformed into cells, the retron system yielded efficiencies of >20%. As a control, we created a plasmid that contained only the gRNA and no retron donor sequence; SSTR events were eliminated and there were very few events with this control plasmid that resulted in lysine prototrophy (Fig. 3B and 3C).

**Figure 3.**
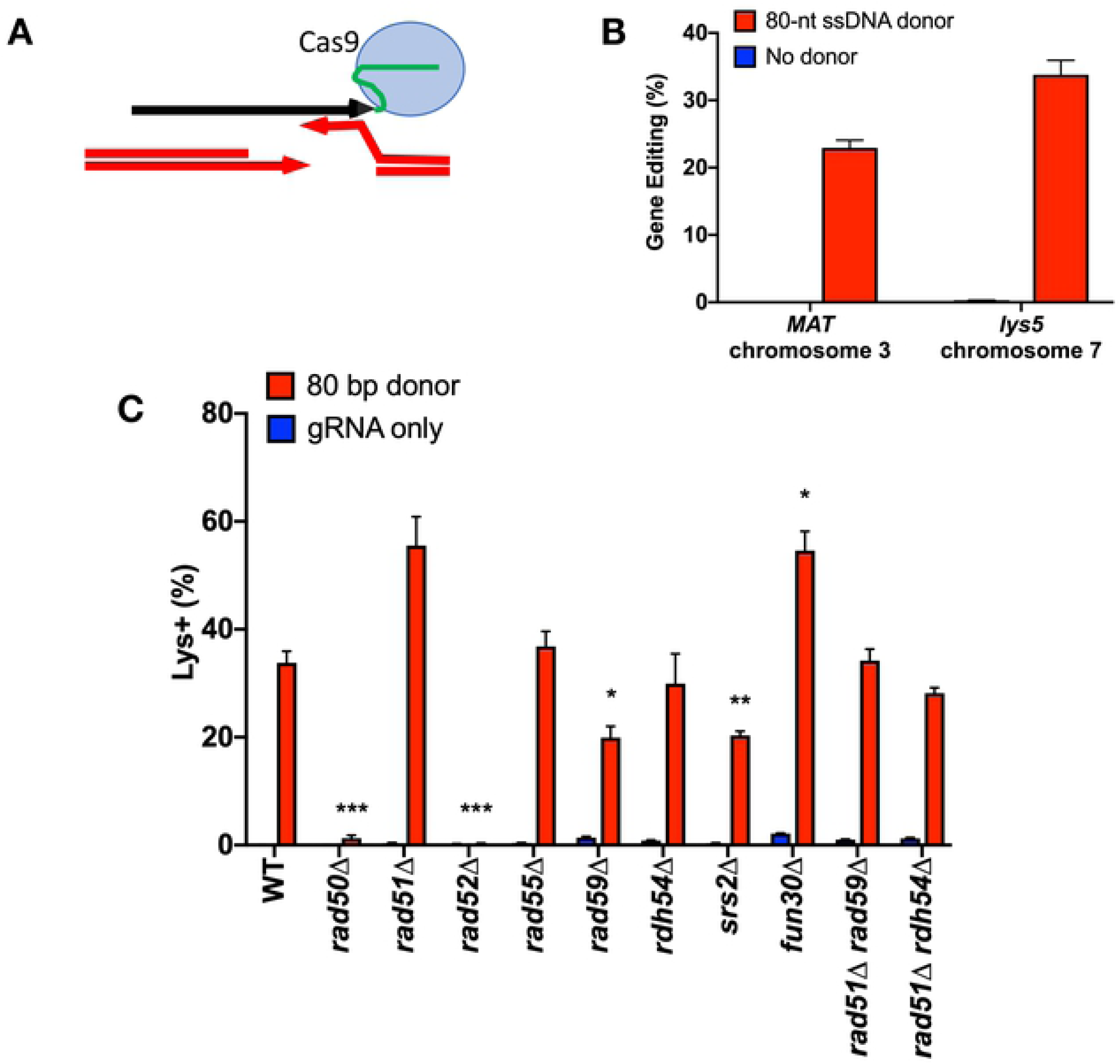
SSTR initiated by retron-Cas9 utilizes a Rad51-indepndent repair pathway. A) The retron system utilizes a modified Cas9 gRNA that tethers the ssDNA donor template to the RNA scaffolding of the Cas9 protein. Successful SSTR at the *MAT* locus results in the insertion of an *Xho*I restriction site, as in Figure 1. SSTR at the *lys5* locus repairs a 5-bp insertion in the *lys5* locus, resulting in Lys^+^ recombinants. B) Efficiency of the Retron-Cas9 system at two chromosomal locations. Cells were plated onto URA^-^ with dextrose (non-induction) and URA^-^ with galactose (induction) media. At *MAT*, % gene editing was determined by PCR and *Xho*I digest of induction survivors. At *lys5*, the percentage of gene editing was determined by replica plating URA-Gal survivors onto Lys^-^ media. The resulting plate count over plate counts on non-induction media results in % gene editing. C) Effect of recombination mutants on retron-Cas9 SSTR gene editing. After induction of the retron system on galactose-containing media, survivors were replica plates to Lys^-^ media. Percentage of Lys^+^ colonies was calculated as a percentage of total cells plated. Significance determined using unpaired t-tests compared to WT, using the two-stage Benjamini, Krieger, and Yekutieli false discovery rate approach, * p ≤ 0.01, ** p ≤ 0.001, *** p ≤ 0.0001

To test whether the genetic components of SSTR were the same with a Cas9-induced DSB, we recreated gene deletions in this strain background. We found that the requirements were the same as for HO-induced SSTR events, being independent of Rad55, Rdh54 and Rad51, but still dependent on Rad52, Rad59, Srs2, and the MRX complex and inhibited by Fun30 (Fig. 3C). With the retron system, the double mutants *rad51*Δ *rad59*Δ and *rad51*Δ *rdh54*Δ do not show the same significant increase in SSTR using the Cas9 endonuclease as observed with the HO-endonuclease. This difference could be due to several different reasons. First, the 5’ end of the retron ssDNA is covalently linked to the gRNA, so the template itself may limit access to the repair machinery from the 5’ end of the template. In addition, the tethering of the template to Cas9 will change the dynamics of the homology search. There could be other differences as well, as Cas9 may stay bound to DNA after cleavage, although how long it remains bound *in vivo* is not clear [63, 64]. Previous results have also shown that Rdh54 can translocate on duplexed DNA and disrupt joint molecules [65]. Continued binding of the Cas9 might prevent this disruption.

### Genetic Requirements of SSTR Depend on Template Design

To test if changing the design of the ssODN donor altered the genetic requirements of SSTR, we used a ssODN similar to that described in Fig. 1A, except that the 37 nt of homology on each side of the *Xho*I site were each targeted to sequences that are 500 bp from the DSB; thus, successful gene editing via the 80-nt ssODN creates a 1-kb deletion flanking the *Xho*I site (Fig. 4A). Successful SSTR should then only occur after extensive 5’ to 3’ resection of the DSB ends. Repair efficiency, as measured by viability, using this donor template was significantly lower than the ssODN that simply incorporated an *Xho*I restriction site (approximately 1% compared to 3%) (Fig. S2).

**Figure 4.**
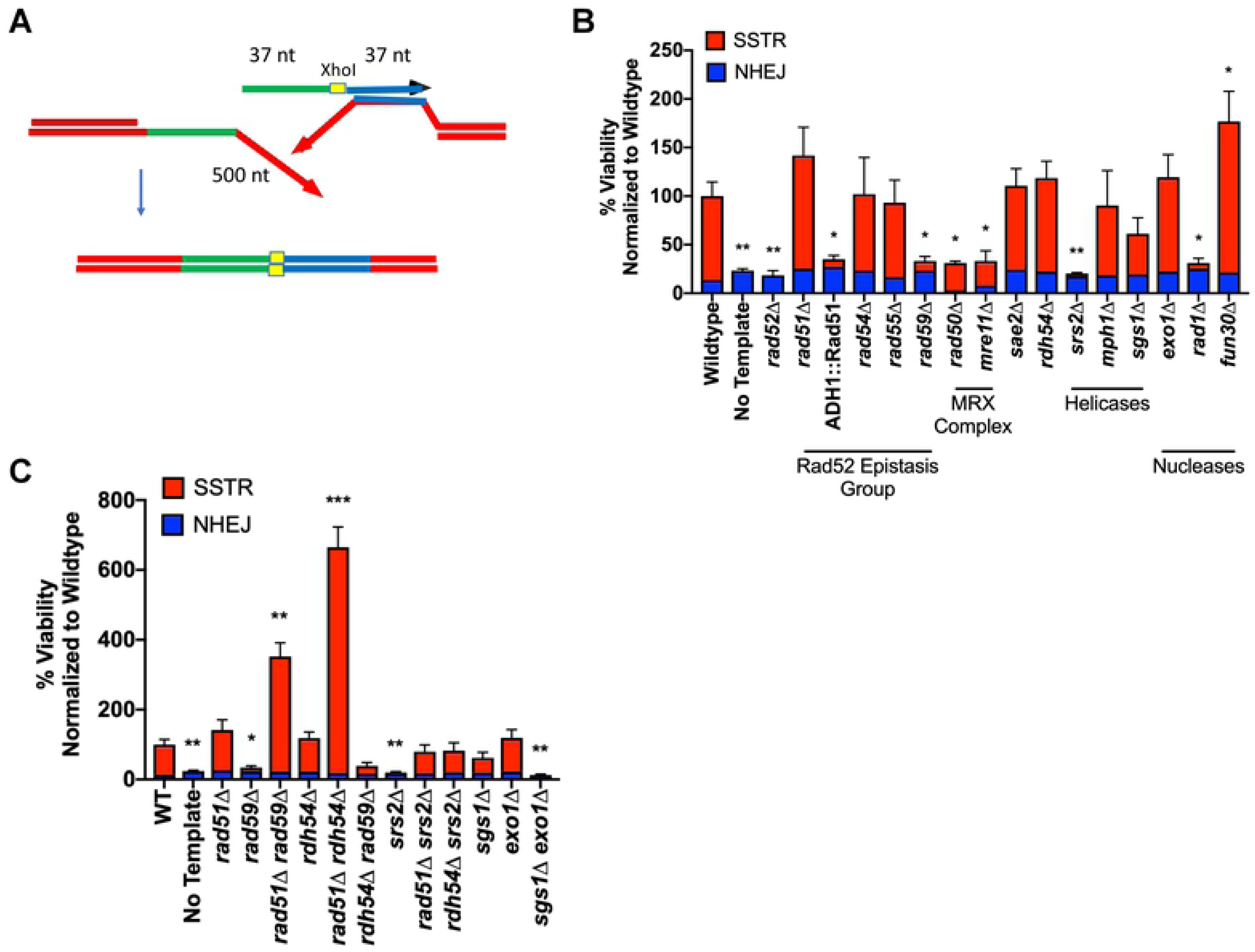
The genetic requirements of SSTR are dependent upon ssODN template design. A) SSTR using a ssODN with 37-nt homologies located 500 bp on either side of the HO-induced DSB. SSTR results in a 1-kb deletion and the incorporation of the *Xho*I restriction site into the *MAT* locus, which can be screened via PCR and restriction digest with *Xho*I. B) Viability determined by galactose-induction plate counts over YEPD plate counts. Proportion of SSTR and NHEJ events in various recombination mutants determined via PCR. C) Effect of double mutants on deletions created by SSTR. Significance determined using a paired t-test with Welch’s correction, * p ≤ 0.01, ** p ≤ 0.001, *** p ≤ 0.0001

The core recombination requirements of SSTR for this configuration were the same as those seen with the simple *Xho*I insertion, as gene editing was independent of Rad51, Rad54, Rad55 and Rdh54, but still dependent on Rad52, Rad59, Srs2, and the MRX complex (Fig. 4B). As before, *srs2*Δ was suppressed by both *rad51*Δ and *rdh54*Δ, and there were still substantial increases in *rad51*Δ *rad59*Δ and *rad51*Δ *rdh54*Δ compared to wildtype or *rad51*Δ (Fig. 4B and 4C). However, using an ssODN that creates a large deletion imposes additional requirements. For the ssODN to pair with a resected DSB end, there must be extensive 5’ to 3’ resection. While deleting either the Sgs1 or Exo1 individually had no significant impact, the double mutant *sgs1*Δ *exo1*Δ abolished SSTR (Fig. 4C). This result stands in contrast to the lack of effect of the double mutant in the simple incorporation of the *Xho*I site and emphasizes the need for long-range 5’ to 3’ resection (Fig. 1C). Also, as above, Sae2 is not required. The effect of blocking long-range resection in *sgs1*Δ *exo1*Δ was not mimicked by deleting Fun30, whereas in previous studies examining resection of DSB ends fun30Δ showed 5’ to 3’ resection of a dsDNA end similar to *sgs1*Δ *exo1*Δ [46, 47, 48].

Another requirement in the deletion assay is for Rad1, and presumably Rad10, which together act as a 3’ flap endonuclease that can remove the 3’-ended 500-nt nonhomologous tail that would be created by annealing the ssODN to its complementary strand [16, 66] (Fig. 4A). Rad1 is not needed in the simple *XhoI* insertion assay (Fig. 1B).

### Single Strand Template Repair Requires both Annealing and Synthesis Steps

How SSTR occurs is still not fully understood. One question concerns the fate of the ssDNA template strand itself. Another concerns the fate of mismatches between the template strand and the complementary single-stranded DSB end. We used the same ssODN described in Fig. 1, but now using ssODNs that contained mismatches in the donor sequence. One donor had 4 mismatches 5’ to the *Xho*I site, while the second had 4 heterologies on the 3’ side, spaced every 9-nt (Fig. 5A). We note that mismatch position 1 is only 2-nt away from the end of the ssODN. We observed that there was a significant decrease in gene editing with 4 mismatches on either the 5’ or the 3’ side, compared to the fully homologous template (Fig. S4) although the effect was more pronounced on the 3’ side (Fig. 5B). This difference may reflect the fact that the initial annealing steps in SSTR can only happen on the side 3’ to the *Xho*I site and may be quite different from the annealing of the second end.

**Figure 5.**
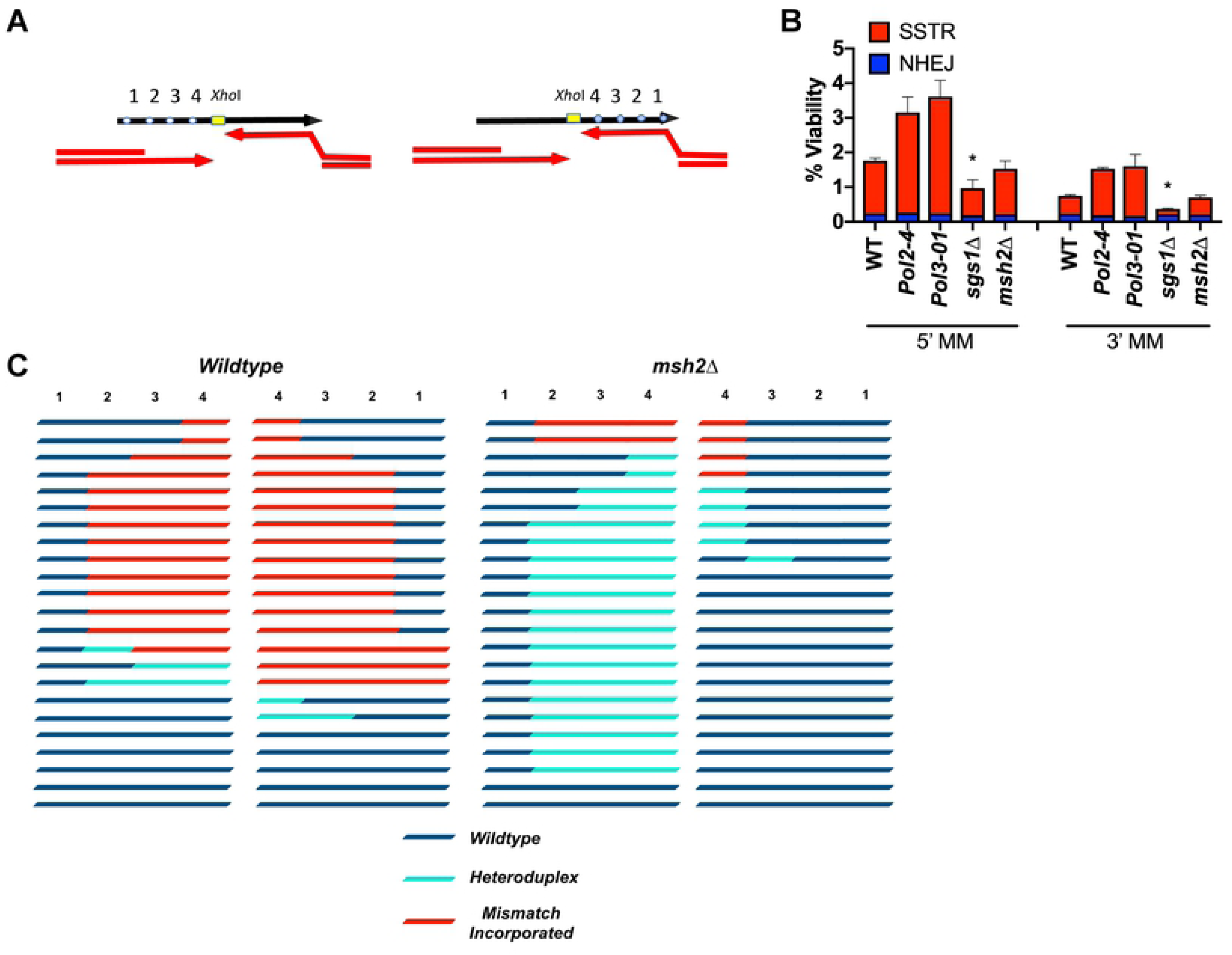
Gene editing events are dependent on the activity of MSH2. A) Experimental set-up to determine the effect of mismatches on SSTR. Mismatches are spaced every 9 nt from the *Xho*I site in the center of the template. Mismatch 1 is located only 2 nt from the terminus of the ssODN. SSTR events are determined by incorporation of an *Xho*I restriction site. B) Significance determined using a paired t-test compared to WT of the same template, * p ≤ 0.01. C) Inheritance of mismatches in wild-type (left) and *msh2Δ* (right) strains. The inheritance of markers was determined separately by sequencing SSTR survivors using ssODNs with mismatches 5’ or 3’ of the *Xho*I site. The different outcomes were grouped. n=23 per template per genetic background.

In studies of SSA and DSB-mediated gene conversion, the inhibitory effects of a small percentage of mismatches could be overcome by deleting Sgs1 or components of the mismatch repair system [67, 68]. Here, however, deleting Sgs1 did not suppress the effect of the four mismatches. In fact, with mismatches on the 3’ side of the *Xho*I site, the majority of survivors in *sgs1*Δ were NHEJ events. Deleting Sgs1 also consistently reduced SSTR in the fully homologous case, though not statistically significantly in any one assay (Fig. 1 and 4).

Deleting the mismatch repair gene, *MSH2*, did not suppress the reduced level of SSTR in the templates carrying 4 mismatches (Fig. 5B), but there was a notable change in the inheritance of these mismatches (Fig. 5C). We analyzed the DNA sequences of 23 SSTR events in both wild type and *msh2*Δ strains for each of the ssODNs carrying mismatches (Fig. 5C). In wild type strains, a majority of *Xho*I insertions were accompanied by co-inheritance of 3 of the 4 mismatches. However, in an *msh2*Δ strain, the majority of SSTR events using the 5’ mismatch ssODN template showed heteroduplex tracts, as evidenced by the presence of both the chromosomal and mutant alleles at these sites when DNA from single transformants were sequenced. On the 3’ side, *msh2*Δ eliminated the great majority of events in which the mutations in the ssODN were inherited into the gene-edited product. These results extend the conclusions reached by Harmsen *et al*. studying SSTR in mammalian cells, where the absence of mismatch repair largely prevented incorporation of heterologies on the 3’ half of the ssODN, while not preventing their assimilation in the 5’ half [69]. Of course in their studies of mammalian cells, it was not possible to detect the presence of unrepaired heteroduplex DNA, as we show in Fig. 5. Assuming that the ssODN itself serves as a template and is not itself incorporated into the gene-edited product, any incorporation of mismatches on the 3’ end of the *Xho*I site should occur only during the time that the resected DSB end has paired with the donor, to prime DNA polymerase to copy the template strand (Fig. 6). On the 5’ side, however, the initial copying of the template and its subsequent annealing to the second DSB end should obligately produce heteroduplex DNA that will be resolved by mismatch repair to be fully mutant or fully wild type (Fig. 6).

**Figure 6.**
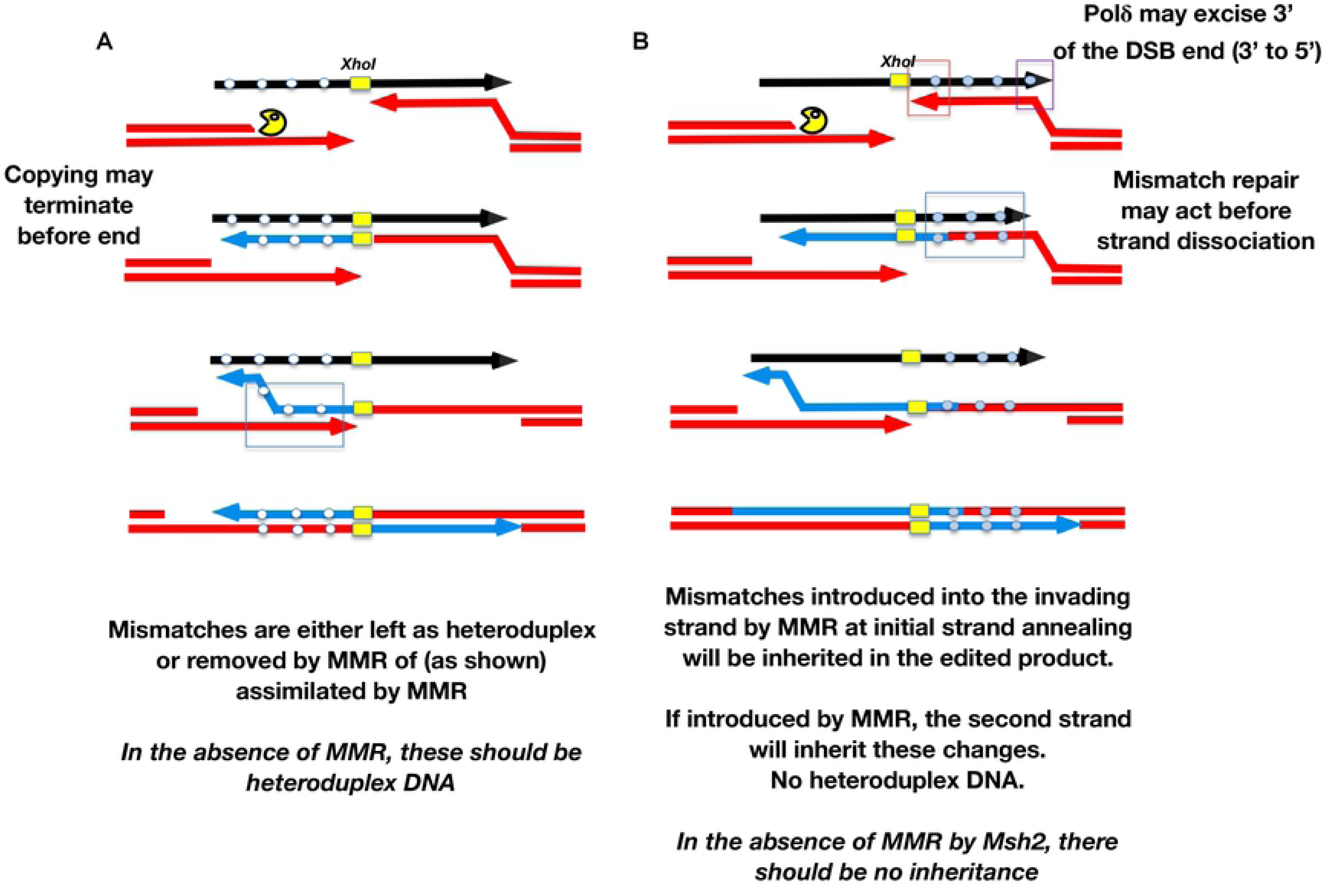
Model of heteroduplex formation and incorporation of mismatches during SSTR. Fate of an ssODN with 4 mismatches either 5’ to the *Xho*I site (A) or 3’ (B). After the DSB is created and resected only the strand on the right can anneal to the template. This annealing creates a heteroduplex that may be repaired by the mismatch repair machinery including Msh2. Heterologies close to the 3’ end of the invading strand, but also at the 3’ end of the ssODN, can be excised by the 3’ to 5’ exonuclease activity of DNA polymerase δ. Only if the heteroduplex is converted to the template strand genotype will these mismatches be incorporated into the SSTR product (B). Mismatches 5’ to the *Xho*I site will be obligately copied by DNA polymerase after strand invasion (A). The dissociation of the newly copied strand allows it to anneal with the resected second end of the DSB, creating an obligate heteroduplex. Dissociation of the newly copied strand may occur without copying the end of the ssODN template. Heteroduplex DNA may then be corrected to the genotype of the donor template or left as unrepaired heteroduplex. In the absence of Msh2, most outcomes will have heteroduplex to the left of the *Xho*I site but no incorporation to the right.

We noted that mismatches located 2-nt from either end of the ssODN were only rarely assimilated into the gene-edited product (Fig. 5C). In our recent study of break-induced replication (BIR), we discovered that heteroduplex DNA created by strand invasion was corrected (i.e. mutations were assimilated into the BIR product) in a strongly polar fashion from the 3’ invading end [70]. Moreover, these corrections of the heteroduplex were orchestrated by the 3’ to 5’ exonuclease (proofreading) activity of DNA polymerase *δ*, which removed up to 40 nt from the invading end and replaced them by copying the template. Incorporations of the mismatched base from the template was almost completely abrogated by eliminating the proofreading activity of DNA polymerase *δ* (*pol3-01*). Here, using the *Xho*I insertion ssODN with 4 mismatches on one side of the insertion or the other, we found that the overall-incorporation of mismatches was unaffected by proofreading-defective mutations in Pol*ε* (*pol2-4*) or Pol*ζ* (*pol3-01*), with one notable exception: the mutation 2-nt from the 3’ end of the template was incorporated at a very high level in *pol3-01* mutants (Fig. 7A). These data suggest that Pol*ζ* can be loaded not only onto the 3’ end of the chromosomal DSB, but can also be recruited by the 3’ DNA end of the ssODN – and then chew back the 3’ end before it might begin synthesizing new DNA. This observation raises the possibility that the ssODN itself could be incorporated into the gene-edited product in some cases. The *pol3-01* mutation did not affect the assimilation of the most terminal mismatch on the 5’ end of the ssODN (Fig. 7A). We suggest that the failure to incorporate this marker into the final product might occur if the DNA polymerase copying the template dissociates before it has copied the last several nucleotides, so this site is not incorporated as heteroduplex DNA involving the second DSB end; alternatively, there could be a Msh2- and DNA polymerase proofreading-independent mechanism that corrects the heteroduplex in favor of the chromosomal sequence (Fig. 6).

**Figure 7.**
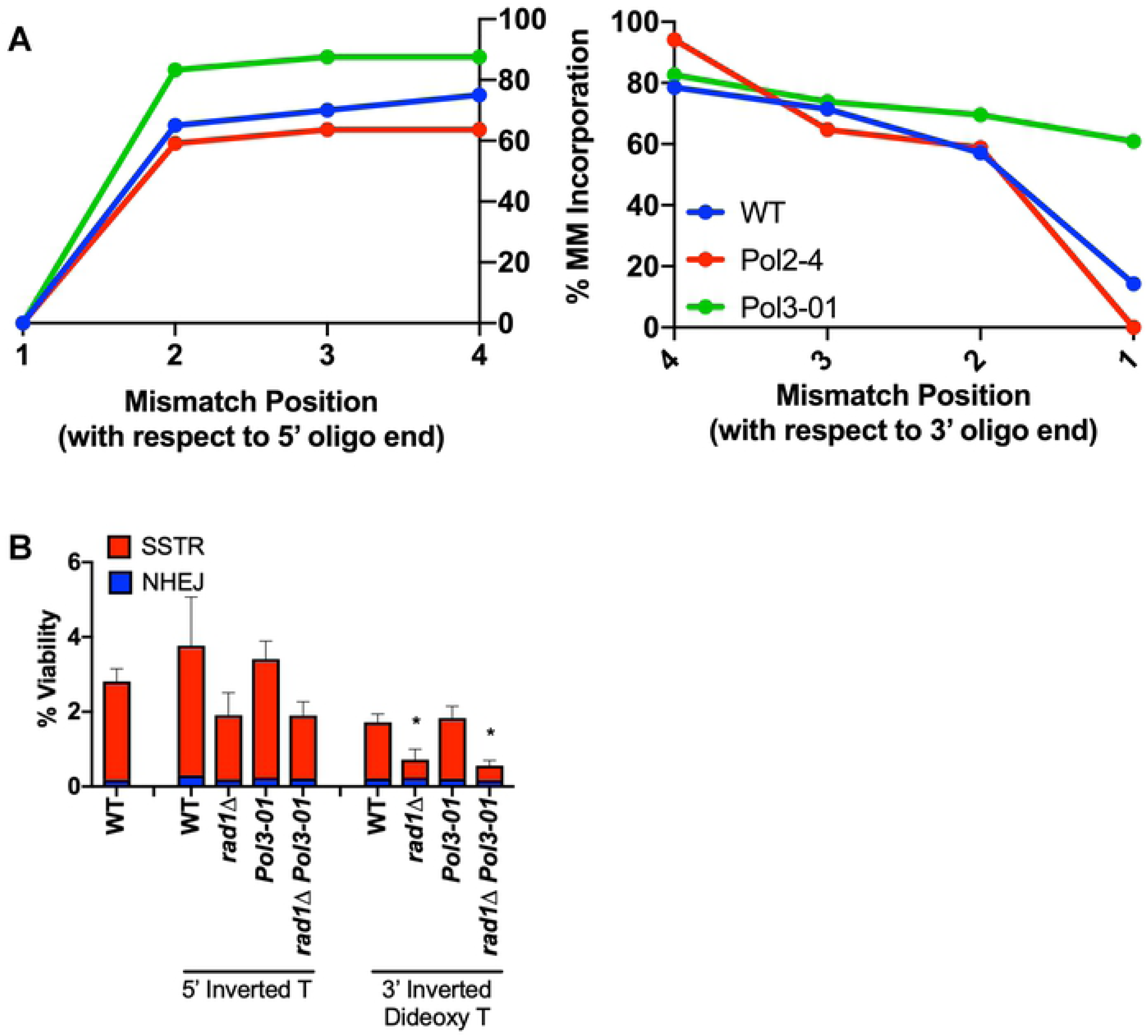
ssDNA templates can be copied or ligated into the DNA to repair DSBs. A) Sequencing of individual SSTR events, n=23. B) Viability test of cells ability to survive DSB repair via SSTR with a template containing a chemical modification on the terminus of the ssODN. Significance determined using a paired t-test with wild-type levels using a template that does not have chemically modified ends, * p ≤ 0.01.

Work by Harmsen *et al*. in mammalian cells also showed that blocking the ends of the template strand reduced SSTR, suggesting that the ssDNA strand itself might be more than a simple template that anneals with a DSB end and is then copied by a DNA polymerase [69]. If the ssODN might be assimilated into the product, then blocking access to either 5’ or 3’ end of the template might affect its usage. We used the *Xho*I insertion ssODN with complete homology to the chromosomal site, except these templates were chemically modified with either an inverted thymine at the 5’ end, or an inverted dideoxy-thymine on the 3’ end. These chemical modifications should prevent ligation of the ssODN into chromosomal DNA, as well as block extension of the 3’ end by Pol*ζ* unless the block is excised. There was no significant difference between either donor template in a wildtype cells or in a *pol3-01* strain compared to the *Xho*I donor template with no chemical modifications. An alternative way that a modified 3’ end nucleotide might be removed would be through the use of the Rad1-Rad10 flap endonuclease. Indeed, we found a modest reduction in SSTR with the 3’ block in a *rad1*Δ strain when compared to wildtype cells (Fig. 7B). These results indicate that the template might sometimes be ligated into the repaired product, or that the inverted thymine causes increased rejection of the ssODN as a suitable repair template.

### SSTR is distinct from dsDNA template repair

Our SSTR assays employ templates containing only 37-nt of homology on either side of the DSB. We wanted to know if this noncanonical repair pathway is specific to ssDNA, or might also apply to repair with dsDNA templates with the same limited homology. We annealed complementary 80-nt ssDNA oligonucleotides to create a dsDNA template with free ends that had 37-bp of perfect homology flanking each side of a 6-bp *Xho*I restriction site. After duplexing, the pool of dsDNA template was treated with S1 nuclease to degrade any remaining non-duplexed ssDNA. We transformed the template into cells using the same protocol used with the ssDNA templates (Fig. 8A). Double-stranded template repair (DSTR) proved to be more than five times as efficient as SSTR (Fig. 8B). With this short dsDNA template, the repair process shifted to a Rad51-dependent event, now also requiring Rad54, Rad55 and Rdh54, but still dependent on the MRX complex, Srs2 and Rad59 (Fig. 8C).

**Figure 8.**
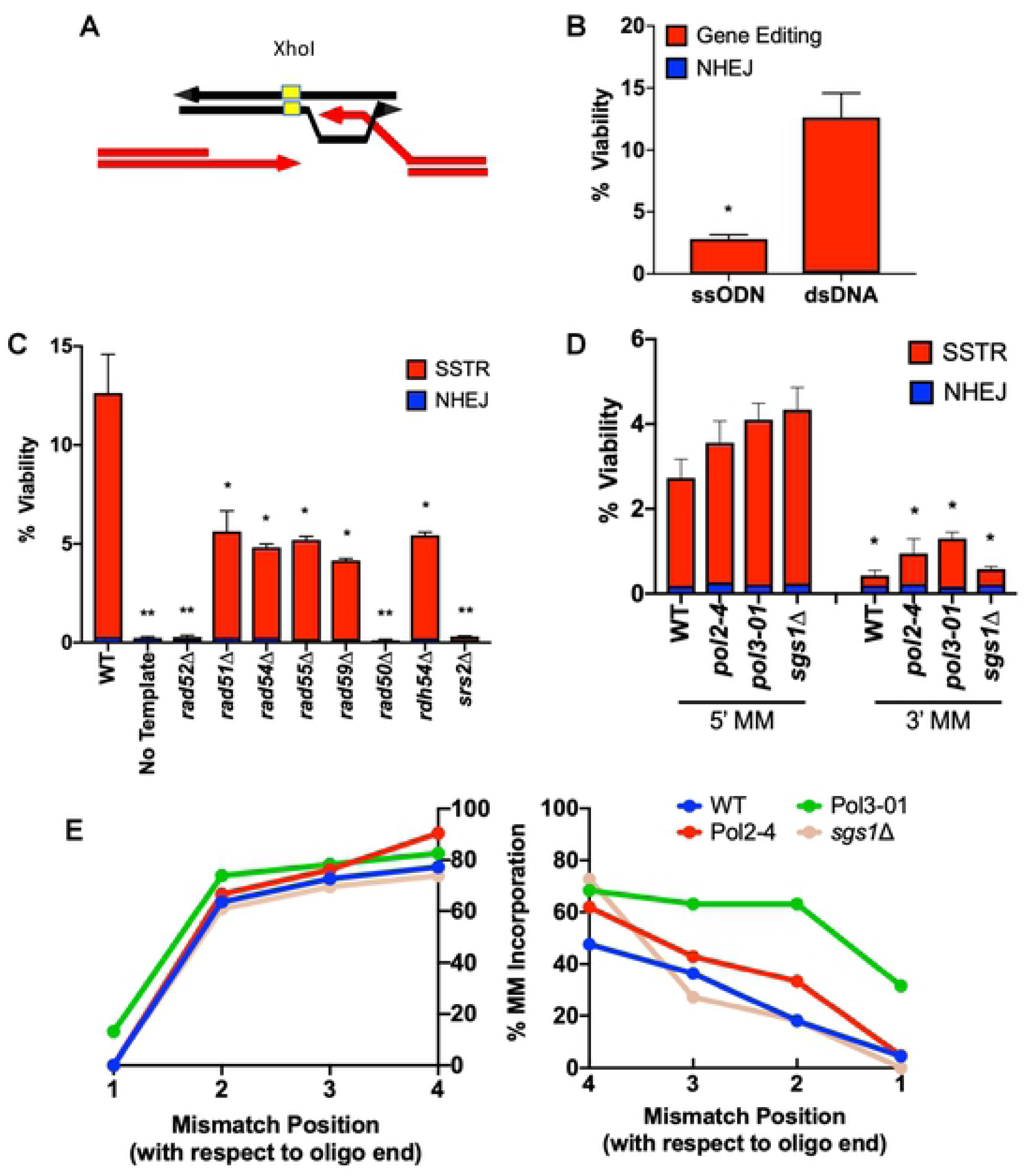
SSTR is distinct from DSTR. A) Successful DSTR results in the incorporation of the *Xho*I restriction site into the *MAT* locus, which is screened for via PCR followed by an *Xho*I restriction digest. B, C, D) Viability tests to determine strains ability to undergo DSB repair with designate 80-nt ssDNA or 80-bp dsDNA template. Viability determined via plate counts of galactose-induction media over YEPD non-induction media. Significance determined using a paired t-test with Welch’s correction, * p ≤ 0.01, ** p ≤ 0.001. E) Sequencing of *MAT* locus following repair from each of two 80-bp templates, with 4 mismatches at the 5’ or 3’ side of the *Xho*I site (n=24 in each case).

With DSTR templates carrying 4 mismatches on either side, the efficiency of repair dropped markedly compared to wildtype, similar to what we see with SSTR (Fig. 8D). We note that because both the resected 3’ end of the DSB on the left or the right could initiate strand invasion, and the template itself is double-stranded, we cannot account for the difference in the effect of the mismatches on the upstream or the downstream side in the same terms we used with the SSTR substrates. It is possible that the two ends of the DSB do not participate equally in initiating DSTR. This possibility is supported by sequencing data, which show that in wildtype cells a mismatch 9 bp away from the upstream side of the DSB will be incorporated into the genome in 78% of survivors, while a mismatch 9 bp away from the downstream side of the DSB will only be incorporated in 48% of survivors (Fig. 8E). Additionally, the *pol3-01* proofreading-defective allele of Pol*ζ* shows a higher rate of mismatch incorporation on the downstream side of the DSB compared to wildtype cells, indicating that Pol*ζ* may still be resecting and replacing the 3’ end of the annealed template strand. Given the apparent symmetry of this DSTR system, it is difficult to explain the different results on the upstream and the downstream sides of the template.

## Discussion

Although CRISPR/Cas9 has made it possible to generate specific changes to the genome in many organisms, the model organism of budding yeast still serves as an important platform to determine the mechanism of gene editing, and thus optimize experimental design. Here we have studied targeted gene editing events after a DSB and made several important observations.

Gene editing using ssODN templates utilizes a novel pathway of DSB repair that is independent of the canonical repair proteins Rad51, Rad54, Rad55 and Rdh54/Tid1, but still dependent on Rad52, Srs2, and the MRX/MRN complex. When the ssODN templates are designed to create a large deletion, Rad1-Rad10 and the long-range resection machinery also become essential. We have confirmed that the genetic requirements of SSTR are generally the same in HO-mediated SSTR as with Cas9-mediated events. Moreover, the mechanism of SSTR is specific to ssDNA templates, as gene editing using a dsDNA template of the same size and sequence (with 37 bp homology to each DSB end) switches to a Rad51-dependent mechanism that is nevertheless distinct from that involving gene conversion between long regions of homology.

We have also determined an important role of the Rad52 paralog Rad59, which apparently acts to alleviate the inhibition of Rad51 on Rad52’s ability to anneal ssDNA tails. Previous studies have shown that Rad51 impairs Rad52’s ability to anneal DNA strands, but this can be overcome by Rad59 [43]. In SSTR, deleting Rad51 suppresses *rad59Δ*. This suppression is not only true in SSTR, but in the more canonical DSB repair pathway of SSA. We note that suppression of *rad59Δ* by *rad51Δ* is quite a different relationship than is seen in spontaneous recombination between chromosomal regions, where *rad51Δ rad59Δ* is much more severe than *rad59Δ* or *rad51Δ* alone [71]. In some of our assays the efficiency of SSTR is significantly greater for *rad51Δ rad59Δ* than for *rad51Δ* alone; this result suggests that Rad59 may impair SSTR in other ways than simply modulating Rad51. Rad59 interacts directly with Rad52 and may form heteromeric rings [72, 73]; possibly without Rad59, Rad52’s annealing activity is intrinsically greater. We note that Rad59 is also important in DSTR with short homology, while its role in DSB repair with longer homology is not critical [21].

SSTR shares some features with other HR pathways that involve short regions of homology [22, 25], such as the need for the MRX complex. In DSB repair events that involve longer (>200 bp) homology, deleting components of MRX delays but does not diminish HO-induced DSB repair [74]; yet with short substrates MRX plays a central role, both for SSTR and DSTR. Yeast MRX has been implicated in many early steps in DSB repair [75]; it is required for most NHEJ events, can bridge DSB ends, promote short 3’-ended ssDNA ends by 3’ to 5’ resection, promote the loading of DSB-associated cohesin, and more. How it is implicated in SSTR and DSTR is not yet clear. Similarly, the Srs2 helicase is critical for SSTR (and DSTR). The role of Srs2 as an anti-recombinase, to displace Rad51, appears to account for its importance, as *rad51Δ* suppresses *srs2Δ*; but we do not yet understand why *rdh54Δ* also suppresses *srs2Δ*.

As expected, the presence of mismatches in the ssDNA template reduces the efficiency of SSTR. However, the degree of inhibition for the level of heterology we used – 4 mismatches in a 37-nt region (∼11% divergence) – was only about 4-fold, quite different from the greater than the 700-fold reduction seen between dsDNA inverted repeat substrates [76]. Moreover, we were surprised that the effect of these heterologies was not suppressed by deleting either Msh2 or the helicase Sgs1, as previous studies have shown that recombination between divergent sequences – both gene conversions between dsDNA sequences and single-strand annealing – is markedly improved by deleting Sgs1 and components of mismatch repair [67, 68]. For Sgs1, the 11% level of heterology is greater than that studied in SSA (3%) or spontaneous inverted repeats (9%), so it is possible that Sgs1 cannot respond to such a level of divergence. However, we note that the ability of Sgs1 and Msh2 to discourage recombination of heterologous sequences is apparently quite different when the DSB end contains a nonhomologous tail versus (as in the cases studied here) ends that do not have such sequences [70]. Alternatively, Sgs1 may play a specific role in SSTR that has not yet been revealed.

Additionally, we have provided evidence that incorporation of mismatches templated by the ssODN into the genome occurs in an Msh2-dependent manner. SSTR provides an unambiguous way to distinguish between the initial strand annealing event with the one DSB end that is complementary to the ssODN and the subsequent events that lead to second end capture and the completion of DSB repair. Confronted with 4 mismatches in the ssODN on either side of the DSB, the cell efficiently incorporates these heterologies into the gene-edited product, but by two distinctly different processes. The initial annealing of the resected end with the template produces a heteroduplex DNA that should be short-lived, until the 3’-extended strand dissociates and anneals with the second resected end. Only during this time can mismatch repair transfer the heterology to the strand that will be incorporated into the final gene-edited product (Fig. 6). We have previously shown that such events occur rapidly during an HO-induced gene conversion (*MAT* switching) and depend on mismatch repair machinery [77]. In addition, we find that a mismatch very close to the 3’ end of the ssODN is usually not incorporated, unless the 3’ to 5’ proofreading activity of DNA Polymerase *δ* is eliminated. We demonstrated a similar type of proofreading in BIR, where the 3’ strand invading into a duplex DNA donor is resected [69]. By the same token, 3’ to 5’ resection of the DSB end will assure that a heterology close to that end (close to the *Xho*I site) could be incorporated without the need for Msh2.

Once the annealed end is extended, copying the 5’ side of the ssODN, all of the heterologies on that side will be copied; but when this newly-copied strand anneals with the second DSB end, there will be an obligate heteroduplex. Hence, on this side, in the absence of Msh2, we recover sectored colonies. The failure to incorporate the 5’-most heterology may reflect dissociation of the newly copied strand before it reaches the end of the template.

These considerations lead us to the model of SSTR shown in Fig 6 and the graphical abstract. After a DSB, the cell initiates end resection, forming ssDNA tails. This process may require MRX proteins to create 3’-ended tails. In the absence of this complex it is possible that neither Exo1 nor Sgs1-Rmi1-Top3-Dna2 can act soon enough to permit use of the ssDNA template before its degradation; however, MRX is required even when ssDNA templates are generated by the retron system, which presumably has the capacity to create ssDNA continually while under induction. Strand annealing depends on Rad52 and on the action of Rad59 to thwart a Rad51-mediated inhibition of Rad52’s annealing activity. Pol*δ* is also engaged in its proofreading mode to remove heterologies near either the 3’ end of the DSB or the template. In SSTR, the degree of resection of the ssODN is quite limited, as only a marker 2-nt from the end is affected by this “chewing back”. When strand invasion occurs during BIR, the 3’ to 5’ excision can extend up to about 40 bases [70]. Interestingly, the DSTR Pol*δ* proofreading is also apparently more extensive (Fig. 7). Once the first end is annealed and extended, the newly synthesized DNA must dissociate and anneal to the second end of the DSB, thus bridging both sides of the break and creating heteroduplex DNA that is subject to mismatch repair. When SSTR is used to create a large deletion, long-range resection is required and the Rad1-Rad10 flap endonuclease becomes essential. The final gap-filling is carried out by the translesion polymerase Pol*ζ*.

Finally, we have also determined that SSTR is a highly mutagenic event. Until now, the off target-effects of gene editing have primarily been thought of as a result of non-specific cutting of the endonuclease. Here we found that filling-in of the gaps created by long-range resection machinery following the DSB is a highly mutagenic process, dependent on DNA Pol*ζ*. It is therefore highly possible that genes adjacent to targeted DSBs have been mutated during SSTR. Gene-edited products should be screened for these potential mutations.

## Methods

### Parental Strain

JKM179 (*ho*Δ *MAT*α *hml*Δ::*ADE1 hmr*Δ::*ADE1 ade1-100 leu2-3,112 lys5 trp1::hisG*ʹ *ura3-52 ade3::GAL::HO*) was used as the parental strain in these experiments. This strain lacks the *HML* and *HMR* donor sequences that would allow repair of a DSB at *MAT* by gene conversion, and thus all repair occurs through NHEJ or via the provided ssODN. ORFs were deleted by replacing the target gene with a prototrophic or an antibioitic-resistance marker via the high-efficiency transformation procedure of *S. cerevisiae* with PCR fragments^79^. Point mutations were made via Cas9 and an ssODN template. gRNAs were ligated into a *BplI* digested site in a backbone that contains a constitutively active Cas9 and either an HPH or *LEU2*-marker (bRA89 and bRA90, respectively). Plasmids were verfieid by sequencing (GENEWIZ) and transformed as previously described^80^.

### Retron Plasmid Construction

pZS165, a yeast centromeric plasmid marked with *ura3* for the galactose inducible expression of the retron-guide chimeric RNA with a flanking HH-HDV ribozyme^57^ obtained from addgene. gBlocks were designed that contained a gRNA and donor sequence and were cloned into the *NotI*-digested pZS165 backbone using the NEBuilder HiFi DNA Assembly Cloning kit. Integration was verified by sequencing (GENEWIZ).

### SSTR Viability Using HO endonuclease

Strains were grown overnight in selective media, and were then diluted into 50 mL of YEPD and grown for 3 hours. Cells were then pelleted, washed with dH2O, and resuspended in 0.1M LiAc. After pelleting, 25 *μ*L of 100 *μ*M ssODN, 25 uL of TE, 25 *μ*L of 2 mM salmon sperm DNA, 240 *μ*L of 50% PEG, and 36 *μ*L of 1M LiAc was added to the pellet and vortexed. Reactions were incubated at 30°C for 30 minutes, followed by a 20-min incubation at 42°C. Cells were then diluted 1000-fold and plated onto YEPD and YEP-Galactose (YEP-Gal) media. YEPD plates were grown at 30°C for 2 days, and YEP-Gal plates were incubated at 30°C for 3 days. Colonies were then counted to obtain average viability. SSTR vs NHEJ events were determined by pooling survivors and amplifying the *MAT* locus via PCR. Following amplification, PCR products were digested with the *Xho*I nuclease and quantified via gel electrophoresis.

### SSTR Viability Using Cas9/Retron System

Cas9 and the Ec86 retron were integrated into *his3* on chromosome 15 into strain JKM179 that had *ade3::GAL::HO* deleted by restoring *ADE3*. After the plasmid containing the gRNA and Ec86 donor sequence was introduced, strains were resuspended in dH2O and plated onto uracil drop-out media or uracil drop out media containing galactose (URA-Gal). URA plates were incubated at 30°C for 3 days, and URA-Gal plates were incubated at 30°C for 4 days. After incubation, plates were counted to obtain viability. URA-Gal plates were then replica plated to lysine drop-out media to obtain SSTR levels.

### DNA Sequence Analysis

Using primers flanking the region of interest, PCR was used to amplify DNA from surviving colonies. PCR products were purified and Sanger-sequenced by GENEWIZ. The sequences were analyzed using Serial Cloner 2-6-1.

## Acknowledgements

This work has been supported by the National Institute of Health grants R35 GM127029 to J.E.H and GM080600 and GM125650 to G.I. .GM. D.N.G. has been supported by NIGMS Genetics Training Grant T32GM007122 and by the National Science Foundation Graduate Research Fellowship Program under grant 1744555. A.J. was supported by a Research Education for Undergraduates (REU) grant from the National Science Foundation. Any opinions, findings, and conclusions or recommendations expressed in this review are those of the authors and do not necessarily reflect the views of the National Science Foundation.

## Graphical Abstract

1) MRX binds to the ends of the DSB, initiating end resection. 2) Rad51 binds to ssDNA tails. 3) The 3’ end of the ssDNA template anneals to complementary resected DNA, dependent on Rad52. A mismatch in homology is shown in red. 4) A DNA polymerase synthesizes across the template. 5) Rad52 anneals the newly synthesized and displaced DNA to the 5’-3’ resected second end of the DSB. 6) Msh2 corrects heteroduplexes that form, fixing edits into the genome. 7) Second-strand synthesis occurs. 8) Gaps are filled in by translesion Pol*ζ*.

## Supplemental Figure Legends

**<u>S1</u>**<u>. *SSTR can be determined by PCR and restriction digest with XhoI.*</u> The percentage of SSTR after the experiment described in Figure 1 was determined by PCR across the *MAT* locus, followed by restriction digest. NHEJ events will result in a non-digested product, which runs at 2,196 bp, while SSTR results in two bands at 1,829 bp and 467 bp. The intensity of the bands was quantified as shown. Trial 1 and 2 were performed on different days with different sets of media, but with identical protocols.

**<u>S2</u>**<u>. Efficiency of SSTR is different depend on ssODN template design</u> Cell viability following a DSB with transformed 80-nt ssODNs that create a 6-bp insertion versus a 1-kb deletion marked by a 6-bp insertion. Viability determined by colony counts of galactose- induction media over colony counts of YEPD non-induction media. Significance determined using a paired t-test, * p ≤ 0.01

**<u>S3</u>**<u>. *Deletions of translesion polymerases do not affect SSTR efficiency.*</u> Cell viability following and HO-induced DSB with transformed ssODN with 37-nt of perfect homology and a 6-nt *Xho*I restriction site.

**<u>S4</u>**<u>. *Heterologies on either the 5’ or the 3’ end of ssODN affect rate of SSTR.*</u> Cell viability following SSTR with ssODN’s with 4 point mutations on either the 5’ or the 3’ end as the template for repair. Heterologies on either end of the ssODN affect the rate of SSTR, but have a higher affect when clustered on the 3’ end of the donor template. Significance determined using a paired t-test, * p ≤ 0.01

